# Application of Cancer Cell Line Encyclopedia for Measuring Correlation Between Transcriptomics and Proteomics as a Guide for System-level Insights

**DOI:** 10.1101/2024.03.03.583123

**Authors:** Blake Williams, Darryl Perry, Peter Aspesi, Jefferson Parker, Ted Johnson, Wendy Su, Eduardo Tabacman, Kirk Delisle, Kayvon Avishan, Vic Myer, Felipa Mapa, Michael Hinterberg, Alan Williams, Lori Jennings, Nebojsa Janjic, Joseph Loureiro

**Affiliations:** Somalogic, Inc., Boulder CO 80301; Novartis Institutes for BioMedical Research, Inc., Cambridge MA 02139; NullSet Informatics Solutions, Brookline, MA 02445; Chroma Medicine, 201 Brookline Ave., Boston, MA 02215

**Keywords:** proteome profiling, CCLE, proteotypes, orthogonal validation, aptamer:antigen interactions

## Abstract

Robust and reliable proteome measurements provide mechanistic insights in biomedical research. SOMAmer (Slow Off-rate Modified Aptamer) reagents are modified, DNA-based, affinity reagents that measure defined target proteins with reproducibility and accuracy similar to monoclonal antibodies. Applying SOMAmer reagent technology, we developed SomaScan, a clinical proteome profiling platform with capability to measure 7,523 proteoforms for 6,594 human proteins by UniprotID in small sample volumes (e.g., 55μl plasma or serum). We evaluated the platform by profiling the proteome of a panel of well characterized Cell Line Encyclopedia (CCLE) cancer models. Unsupervised machine learning analyses demonstrate the SomaScan assay distinguishing cell lines on the basis of their proteome signatures, and identifying both tissue-specific and oncogenic pathways. The proteome measured by SomaScan correlates with published CCLE transcriptome at a level comparable to other published transcript to proteome studies. Taken together, we demonstrate that the SomaScan platform is a technically reproducible system suitable for biomedical and clinical applications that reliably illuminates underlying biomolecular mechanisms.

## 1. Introduction

### 1.1. Each SOMAmer reagent is akin to a monoclonal antibody sensitive to a range of proteoforms

SomaScan assay measures a predetermined subset of the proteome. The assay is based on modified DNA-based affinity reagents, which are selected from enormous pools of potential sequences (>10^15^) that spontaneously fold into intricate three-dimensional structures. By this method, SOMAmer reagents with precise shape and functional group complementarity to the intended targets can be selected in successive rounds of affinity selection and amplification (SELEX) [1, 2]. To augment the chemical diversity of these DNA libraries, we have employed base modifications that introduce functional groups overrepresented in protein-protein contacts thus creating hybrid affinity reagents with optimized binding properties. To identify SOMAmer reagents used in SomaScan assay, individual purified proteins are used for in vitro selections. These proteins are a collection of commercially available and privately acquired research-grade purified proteins. Each SELEX-input protein is a carefully prepared and validated research grade exemplar of a Uniprot entity. As is the case for the development of Ig-class antibodies, careful follow-up characterization for each reagent against its intended antigen establishes confidence in the reagent performance against its expected endogenous antigen. Here, we seek to characterize the endogenous SOMAmer:protein binding events by measuring correlations between proteomic data measured on SomaScan assay and transcriptomic data from the publicly-available Cancer Cell Line Encyclopedia (CCLE) database, which builds on efforts to understand the relationship between the transcriptome and the proteome (reviewed in [3]).

### 1.2. SomaScan is a pooled, massively parallel proteome profiling assay applying SOMAmer reagents as probes for endogenous proteoforms

The SomaScan platform assembles a pool of SOMAmer reagents, each separately isolated from a large library of initially randomized nucleic acid sequences that has been evolved for high-affinity binding to a defined protein in vitro. [1, 2]. The in vitro SELEX methodology provides a rapid way to identify SOMAmer reagents that are sensitive and specific to the purified protein of interest, and SomaLogic has applied this at scale to develop the 7,596 measurements on the current SomaScan platform [4]. The SomaScan Assay reads out on a custom Agilent microarray platform. Small volumes of blood or other biological sample (<60μl) are probed with the pool of SOMAmer reagents, and those SOMAmers that retain binding to sample are quantified by microarray as Relative Fluorescence Units (RFU). RFU are a continuous variable suitable for comparative abundance analyses across experimental design factors, which are summarized here as log2 transformed values for suitability to parametric analysis. We sought to evaluate SOMAmer performance in relation to the anticipated endogenous antigen abundance, as inferred by steady-state mRNA levels across a panel of well characterized cell lines.

### 1.3. The Cancer Cell Line Encyclopedia (CCLE) as a model system for proteome profiling

Immortalized cell lines are a cornerstone for biomedical research providing an important experimental system to understand human disease. The CCLE is a set of hundreds of cell lines for which extensive multiomic profiling has been generated and shared in the public domain [5, 6]. We have extended that CCLE profiling to a small panel of cell lines for the purpose of evaluating SomaScan measurement of human cell lysates. We use steady-state RNAseq of these cell lines to set expectation for SOMAmer RFU signal distribution across those cell lines. Previous studies have demonstrated that there is a weak, positive correlation between mRNA abundance and protein abundance [7, 8]. This weak correlation is a well-understood consequence of the fact that both mRNA and protein levels in a cell are highly dynamic in a temporal sense resulting from their ongoing synthesis and degradation. Positive correlation is expected since, as central dogma dictates, mRNA is translated into proteins; however, since proteins are subject to post-translational influences, differences in relative rates of synthesis and degradation can easily erode this correlation [8]. Correlations between mRNA and protein levels of about 40% has been observed in multiple studies [8-10]. We reasoned that the CCLE provides a rich resource to evaluate the performance of SOMAmer reagents on the SomaScan assay measuring endogenous proteins in cellular lysates.

## 2. Materials and Methods

### 2.1. Cell line culture and sample preparation

CCLE cell lines are of human origin and the cell lysate preparations are suitable for proteome profiling by SomaScan [5, 6]. Adherent cells are seeded at 10-20% confluence in triplicate T175 flasks and grown for 48hrs. For each replicate, 4 x 1.2 mls of conditioned media is transferred to 1.5 mL microcentrifuge tubes. Tubes were then centrifuged (16,000g, 15 minutes, 4°C) to pellet cell debris and transferred (500 μL/tube) to 1.0 mL 2D barcoded, screw top, storage tubes. Conditioned media samples were frozen in dry ice/ethanol and then stored at -80°C until ready for shipment. For cell lysate preparation, the remaining supernatant was aspirated and cells were washed with cold PBS. Aspirated plates were lysed with cold lysis buffer (Pierce #87787) plus 1x halt protease & phosphatase inhibitor cocktail (Pierce #78441) (2.5mL/flask ; 10ml Lysis buffer/cell line ; 100ul halt/cell line) at 4°C for 15 minutes. Lysate was transferred to 2mL microcentrifuge tubes and centrifuged to pellet cell debris (16,000g, 15 minutes, 4°C). Lysates were transferred to 2mL/ tubes and quantified protein concentration with the Biorad Reagent System (Thermo Fisher #23227). Samples were transfered to 1.0 mL 2D barcoded, screw top, storage tubes (Thermo Scientific # 3741 ; 220-230ul each aliquot) and frozen in dry ice/ethanol and stored them at -80°C until ready for shipment. Suspension cells were grown in T175 flasks for 48 hrs and then gently centrifuged (1000 rpm, 10 minutes, room temp.) to pellet cells. Conditioned media was collected and processed as above, and the remaining cell pellet was washed in cold phosphate buffered saline (PBS) and cell lysate was processed as above.

### 2.2. SomaScan assay data stewardship and technical quality assessment

SomaScan data are provided as summarized RFU values in a proprietary ADAT (.adat) formatted file, a tab delimited text file. SomaDataIO software provided in R is used to convert ADAT formatted data to an R/Biobase class expressionSet object suitable for analysis [11, 12]. SomaDataIO is publically available at https://github.com/SomaLogic/SomaDataIO and CRAN https://cran.r-project.org/web/packages/SomaDataIO/index.html.

Technical quality control analysis was performed using open source arrayQualityMetrics software [13]. Agilent microarray RFU are log2 transformed for analysis and assayed cell lysate samples are run alongside SomaScan platform technical replicates for Buffer and Calibrator sample types. The technical quality of the SomaScan Agilent microarray data was acceptable by arrayQualityMetrics criteria (analysis not shown).

### 2.3. SomaScan in vitro cell lysate data normalization

Given the technical and intrinsic differences among cell lines, a generalized method for microarray data normalization was applied to control for technical variance in the data. Linear transformations according to local and global raw data observations were applied to the data for normalization. Briefly, per sample spike-in hybridization normalizations were applied, and subsequently the data were centered to the global median. These normalization steps are shared with the core SomaScan_v4.1 assay for qualified sample matrix types, including blood plasma and serum, cerebral-spinal fluid, and urine, and aspire to retain maximal biological signal while mitigating technical attributes of the experimental design.

### 2.4. Unsupervised learning of the SomaScan Assay

Principal component analysis (prcomp) and bootstrapped hierarchal clustering were done initially to evaluate the quality of the submitted samples in relation to the control samples in the experiment, and subsequently to begin exploring biological patterns in the data [14, 15] (http://www.sthda.com/english/rpkgs/factoextra). Visualizations, linear modeling, and central dogma analytics were done using R\Bioconductor and Tidyverse software [11, 12] (Tidyverse -https://joss.theoj.org/papers/10.21105/joss.01686#).

The topic modeling unsupervised machine learning approach was applied to SOMAmer RFU measurements of CCLE cell lines. Multinomial topic models were fitted with non-negative matrix factorization using the R package fastTopics [16, 17]. Models were fitted with both 15 and 30 topics, selecting the best converged model each across 10 unique model initializations.

### 2.5. Public CCLE data resource

The Cancer Cell Line Encyclopedia is a resource maintained by the Broad Institute. Transcriptomic data used in this analysis are available at https://sites.broadinstitute.org/ccle/datasets and https://depmap.org/portal/ccle/. The cell lines measured are summarized in Supplemental file 1.

## 3. Results

### 3.1. Preclinical application of SomaScan assay

SomaScan data measures in parallel thousands of human proteins. The RFU measurements are made alongside control reagents to monitor the assay run. Density plots and box plots that summarize each SomaScan sample show the distribution of all or subsets of SOMAmer reagents (Fig. 1). RFU signal distribution for test and positive controls samples, which are technical replicates of calibrator and quality control (QC) sample types, are all distinctive from empty buffer negative controls on the platform (Fig. 1A, 1B). For each SomaScan assay run, control reagents provide a measure of technical performance on the assay (Fig. 1C, 1D). Among the control reagents, we have designed a set of sequences, designated Spuriomers, that have similar composition to SOMAmer reagents, but with sequences that were not evolved to bind to any specific protein in a conventional SELEX experiment. Rather, they are either irrelevant, invented sequences, or reagents that have well-known nucleic acid structural motifs such as stem-loops, G-quadruplexes, pseudoknots and the like (Fig. 1C). Spuriomers provide a measure of non-specific background signal attributed to matrix effects unique to nucleic acid-based affinity reagents. As additional assay controls, hybridization normalization SOMAmer reagents measure spike-in controls for the assay (Fig. 1D).

**Figure 1.**
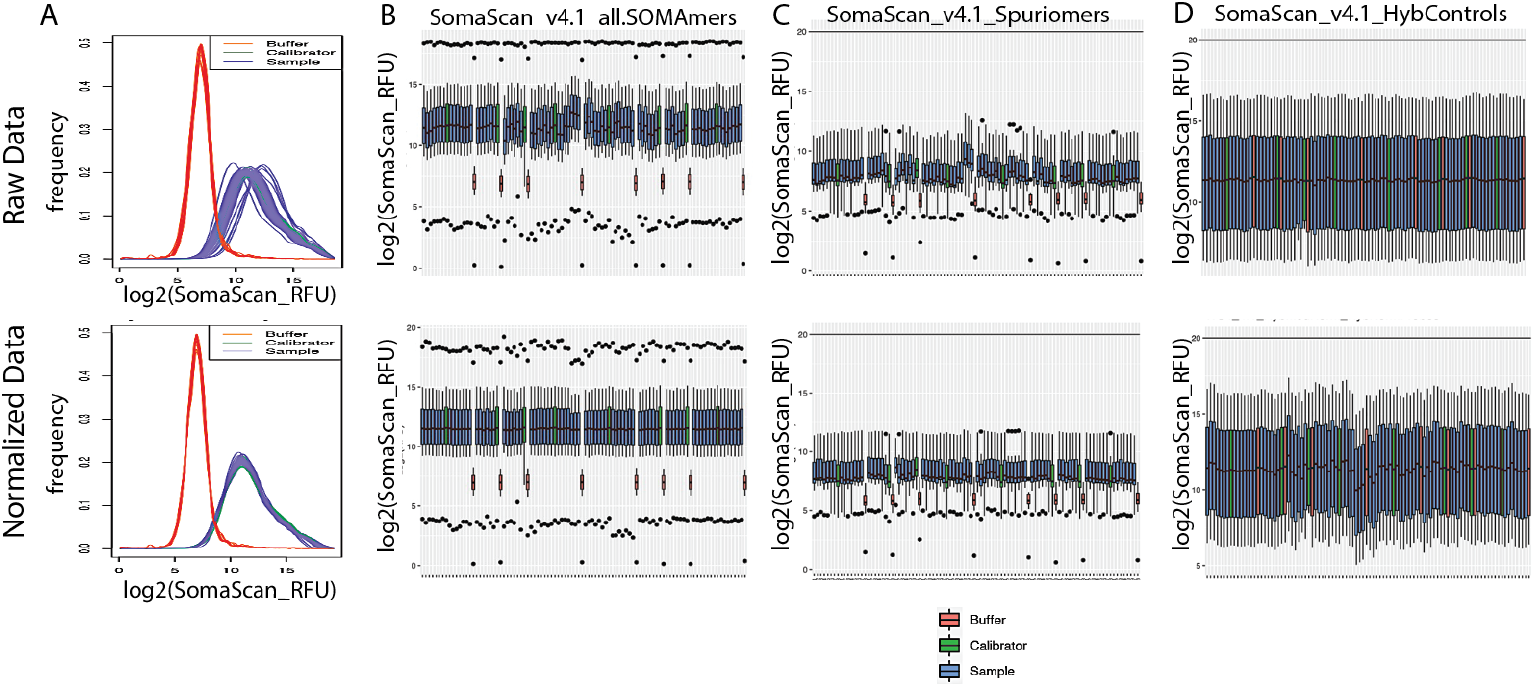
RFU signal distribution for SomaScan assay applied to human immortalized cell culture models. Raw data is shown across the top row and normalized data the bottom row. A) One SomaScan assay per line in the density plots and corresponding box plots in B. B) Box plots of all SOMAmer reagents, C) Spuriomer reagents, and D) Hybridization normalization reagents.

These measurements, in aggregate, integrate biological and technical aspects of the experiment. Unsupervised machine learning methods are apt at uncovering inherent patterns in the data by applying multivariate criteria to measure similarity between samples. We first apply principal component analysis (PCA) to all sample types in the experiment, buffer controls, calibrator controls, and CCLE submitted samples (Fig. 2A, left). In this context we observed expected separation of sample types and low technical replicate variance. When the PCA is done solely on the CCLE submitted samples, a more complex topology is evidence suggestive of cell type co-clustering (Fig2A, right). These patterns are evident in the raw data and are preserved through data normalization.

**Figure 2.**
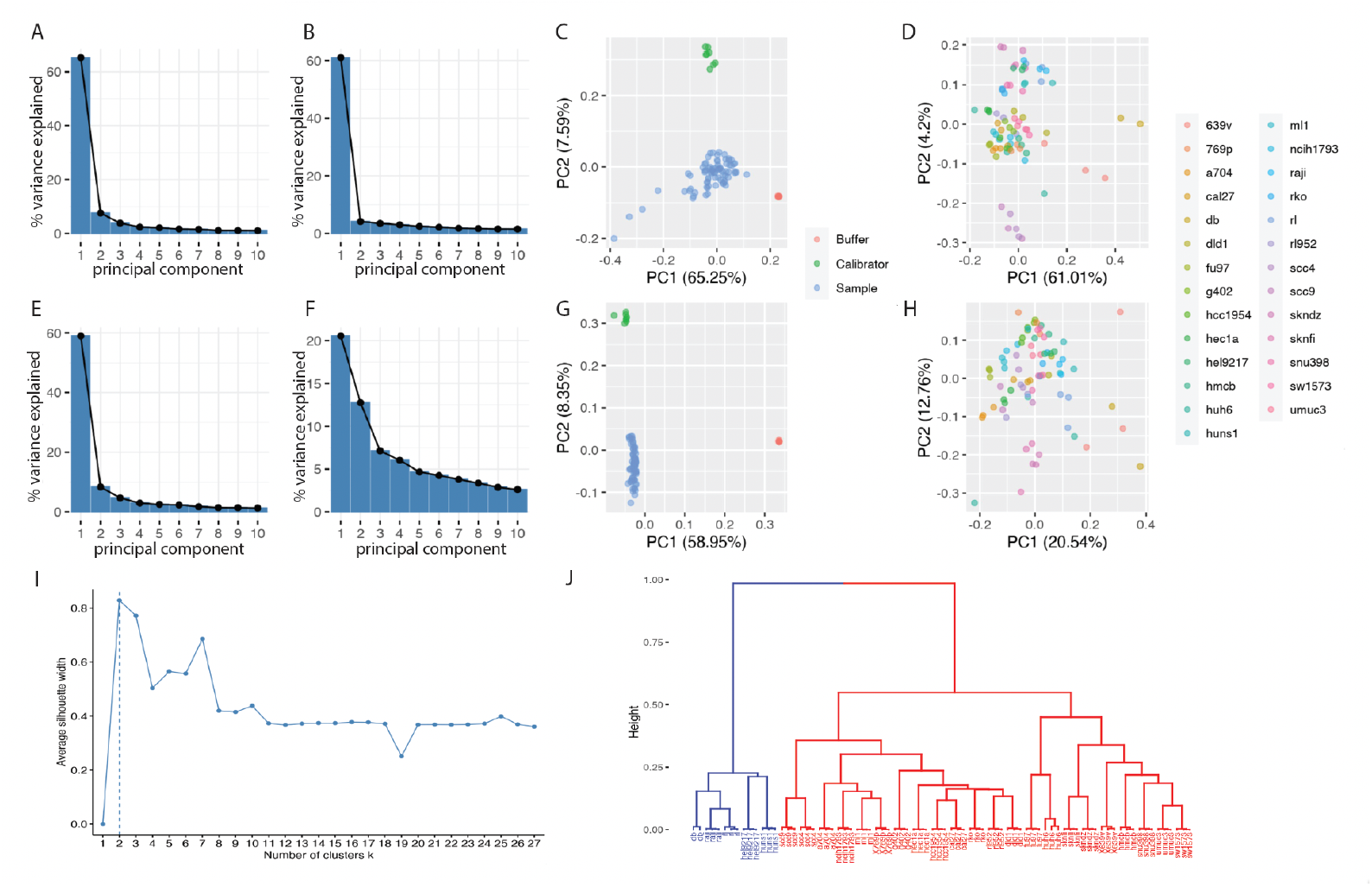
PCA of raw and normalized data. A-D) raw SomaScan Scree plots and PCA for all samples, including technical ones, and separately for the submitted samples. E-H) Normalized SomaScan data Scree plots and PCA, as for raw. E & G) all SomaScan samples, including four replicates of null Buffer and four replicates of standard Calibrator alongside submitted cell lysate samples. F & H) Just the submitted samples from the two plates, which commingle in the first two principal components. I) Unsupervised modeling indicates two distinctive groups, and J) Hierarchal clustering of SomaScan data suggests intrinsic differences between cancer cell lines of hematopoietic versus solid tumor origin, and strong co-clustering of replicate cell lines.

### 3.2. Unsupervised learning by hierarchical clustering identifies distinction between cell lines of solid tumors and those of hematopoietic cancers

Using all SOMAmer reagent measurements as input, we clustered samples according to their multivariate similarity to each other. Hierarchal clustering is a complementary unsupervised method to model the SomaScan data capable of defining sample subsets. Sample metadata are annotated but not used in the derivation of the cluster tree, which identified with confidence two prominent subtypes of samples (Fig 3I). The most prominent distinction in the cell lines belonging to the two subgroups is their tissue origin (Supplemental table 1 – cell line metadata). Notable, hematopoietic cancer cell lines are distinctive from the remaining cancer cell lines of solid tumor origin, indicating that SomaScan is measuring biologically relevant differences among the cancer cell lysate proteomes.

**Fig 3.**
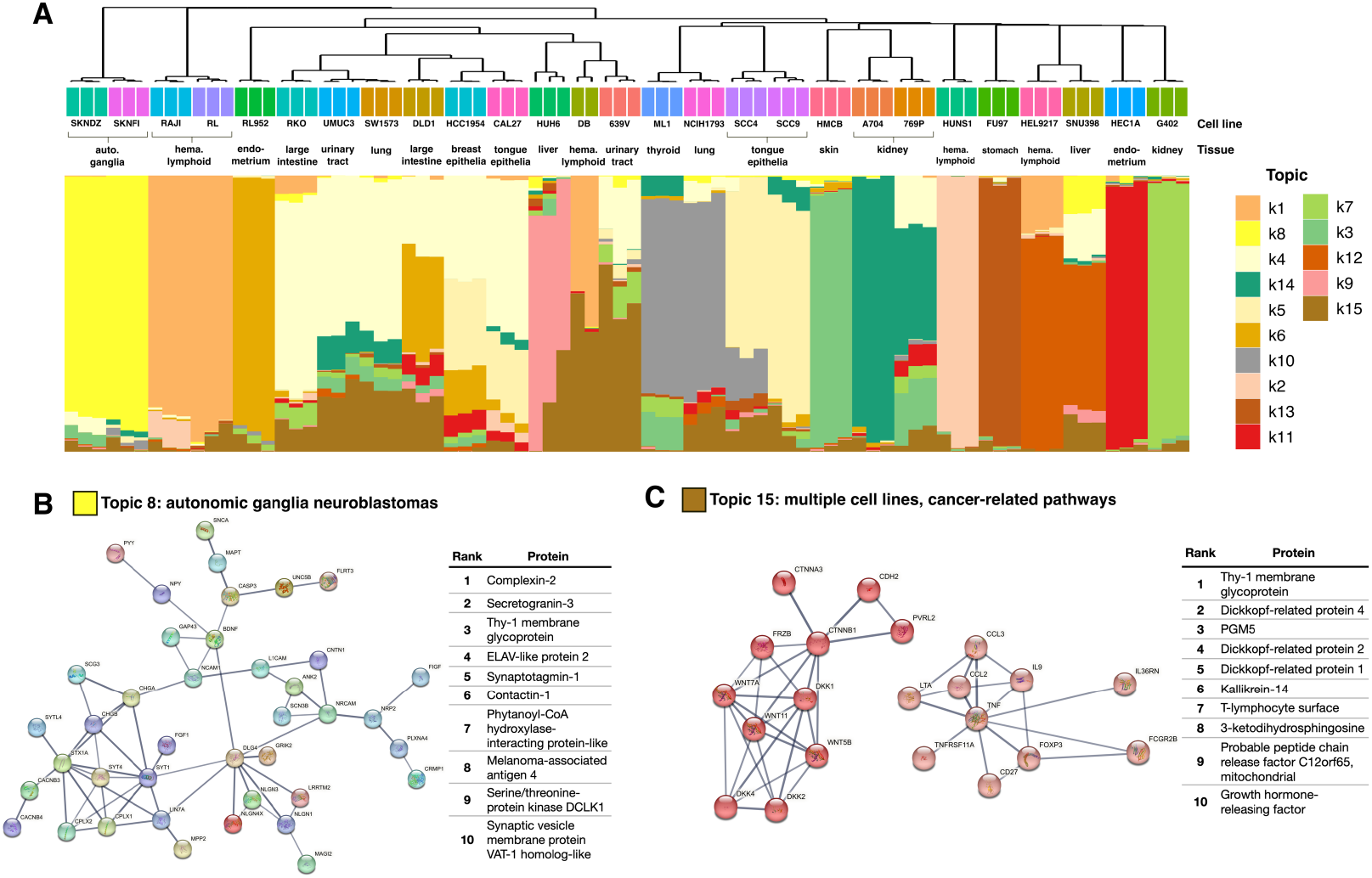
Topic modeling identifies ensembles of target proteins co-modulated in cell line subtypes. A) Samples clustered on topic membership scores reveal clustering by tissue and cancer type. Vertical bars correspond to samples, with the height of each color indicating the relative membership of a sample to each topic. The greater the membership, the more that topic is used to capture protein levels of that sample. Topics are vertically ordered by hierarchical clustering. B) Neuroblastoma-related topic: largest connected component of PPI network of top proteins shown alongside top proteins. C) Multi-cell line cancer-related pathway topic. Key modules of PPI network of top proteins shown alongside top proteins.

### 3.3. Unsupervised learning by topic modeling identifies interpretable dimensions that distinguish cell lines and explain drivers of their proteomes

To demonstrate the richness of proteomic information captured by the SomaScan assay, we applied a more sensitive unsupervised machine learning method referred to as topic modeling to: (i) identify prominent and biologically interpretable dimensions among cancer cell lines, and (ii) use these dimensions to explain, distinguish, and relate cell lines to one another. Although new to proteomic analysis, topic models have been successfully implemented in bulk and single cell RNAseq analyses to identify transcriptomic dimensions [18, 19]. Fit through non-negative matrix factorization, topic models can be thought of as an alternative to principal components analysis, with advantages that include not being constrained to orthogonal dimensions, potentially leading to more biologically coherent dimensions (called topics) [17].

Here we fit multinomial topic models on the SomaScan assay version 4.1 7k panel of RFU values (80 samples: 3 per cell line, except 2 for DB). We first construct a 15-topic model with approximately half as many topics (dimensions) as cell lines to identify topics that span multiple cell lines. This allows us to identify proteomic signals across cell lines related to cancer pathology and tissue type.

In this 15-topic model, samples from the same cell line are clustered together in the space of reduced dimensions defined by the model (Figure 3A). Furthermore, cell lines of similar tissue source and cancer type are more broadly clustered together (e.g., neuroblastoma lines SKNDZ and SKNFI, renal cell adenocarcinoma lines A704 and 769P). Closely clustered cell lines of similar origin tend to have a large membership to the same topic: all samples from SKNDZ and SKNFI are driven by topic 8 (Figure 3B), while A704 and 769P samples are associated with topic 14, and hematopoietic lymphoid lines RAJI and RL are associated with topic 1. Characterization of these topics can describe dominant and shared features of these cell lines. Other cell lines, such as topic 15, explain a portion of the proteome across many samples. These topics may capture cancer-related pathology that is less specific to tissue type.

The aforementioned neuroblastoma-related topic 8 is an example of a tissue-specific topic, defined by proteins indicative of both tissue source (autonomic ganglia) and oncogenic proteins. Enrichment of the 156 top ranked proteins of topic 8 identified tissue-specific pathways: synaptic signaling (fishers exact test, FDR = 7.0 <u>x 10^−9^</u>), neurexin family protein binding (FDR = 1.8E-05), and nervous system development (FDR = 3.7 <u>x 10^−4^</u>), among other enrichments. Top proteins are consistent with this observation, including complexin 2 (synaptic regulator, rank 1 within topic), synaptotagmin 1 (synaptic vesicle membrane protein, 5), and the oncogenic contactin 1 (neuronal surface interaction mediator, 6). Also among the defining proteins of neuroblastoma topic 8 are proteins whose genes are overexpressed with unfavorable neuroblastoma outcomes [20]: neuromodulin (*GAP43*, rank 18), DOPA decarboxylase (*DDC*, 23), and dihydropyrimidinase-related protein 1 (*CRMP1*, 150).

Some topics were tissue non-specific yet still captured cancer-related proteins and pathways (e.g. topic 15). Among the 361 defining proteins of topic 15 are two prominent protein-protein interaction network modules, one capturing WNT signaling and the other capturing tumor necrosis factor and related cytokines (Figure 3C). Within the WNT module is betacatenin, a known proto-oncogene implicated in basal cell carcinoma, squamous cell carcinomas, and other cancers [21]. The highly ranked proteins of topic 15 also include WNT inhibiting Dickkopf-related proteins 4/2/1 (respectively ranked by weight as the 2, 4, 5 top proteins of the topic). Also among the top proteins are regulators of proliferation and differentiation, including growth hormone releasing factor (rank 10), tumor protein 63 (rank 27) and numerous growth factors (topic rank in parentheses): vascular endothelial growth factor D (24), epidermal growth factor-like protein 6 (29), and fibroblast growth factors 22 and 6 (72 and 84, respectively).

A 30-topic model was also fit, since the number of cell lines (27) serves as an approximate expectation for the number of topics that sufficiently explain the RFU matrix (see Supplemental file 2). In this model, every cell line was associated with a single topic (one topic per one cell line) while preserving the perfect within-cell line clustering of the 15 topic model. This allows for a description of each cell line through the analysis of just one topic.

### 3.4. Transcriptome-guided interpretation of SOMAmer reagent RFU signal distribution

Previous work has demonstrated a moderate, positive correlation between mRNA abundance and translated protein abundance in various sample types and organisms by different measurement modalities [8, 22, 23]. We evaluated that relationship between the transcriptome and proteome using SomaScan on CCLE lines (Supplemental file 3). We calculated the mRNA to protein correlation across the 27 cell lines between each SOMAmer reagent RFU and the corresponding mRNA measurements. The resulting distribution of the r^2 over all possible SOMAmer reagents-mRNA pairs, displayed below, has a median value of ∼ 0.10 (Fig. 4A). We recognized that because some transcripts are not expressed in the cell lines profiled, it is not possible to establish a correlation with a protein, as measured by the SOMAmer. We, therefore, focused our attention on the remainder of the measurements where there was measurable mRNA for comparison. Compared to the global correlation of the platform, 2,392 SOMAmers (32.5%) have a compelling positive association (r^2 = 0.371 for those SOMAmers in aggregate) (Fig. 4B), suggesting that for those analytes, the endogenous protein is recognized by the SOMAmer reagent elicited to the SELEX input protein. Furthermore, this overall pattern for those SOMAmers was consistent across the individual cell lines (Fig. 4C). For the cell lines profiled, these reagents have independent corroboration for measurement of the annotated endogenous proteins.

**Fig 4.**
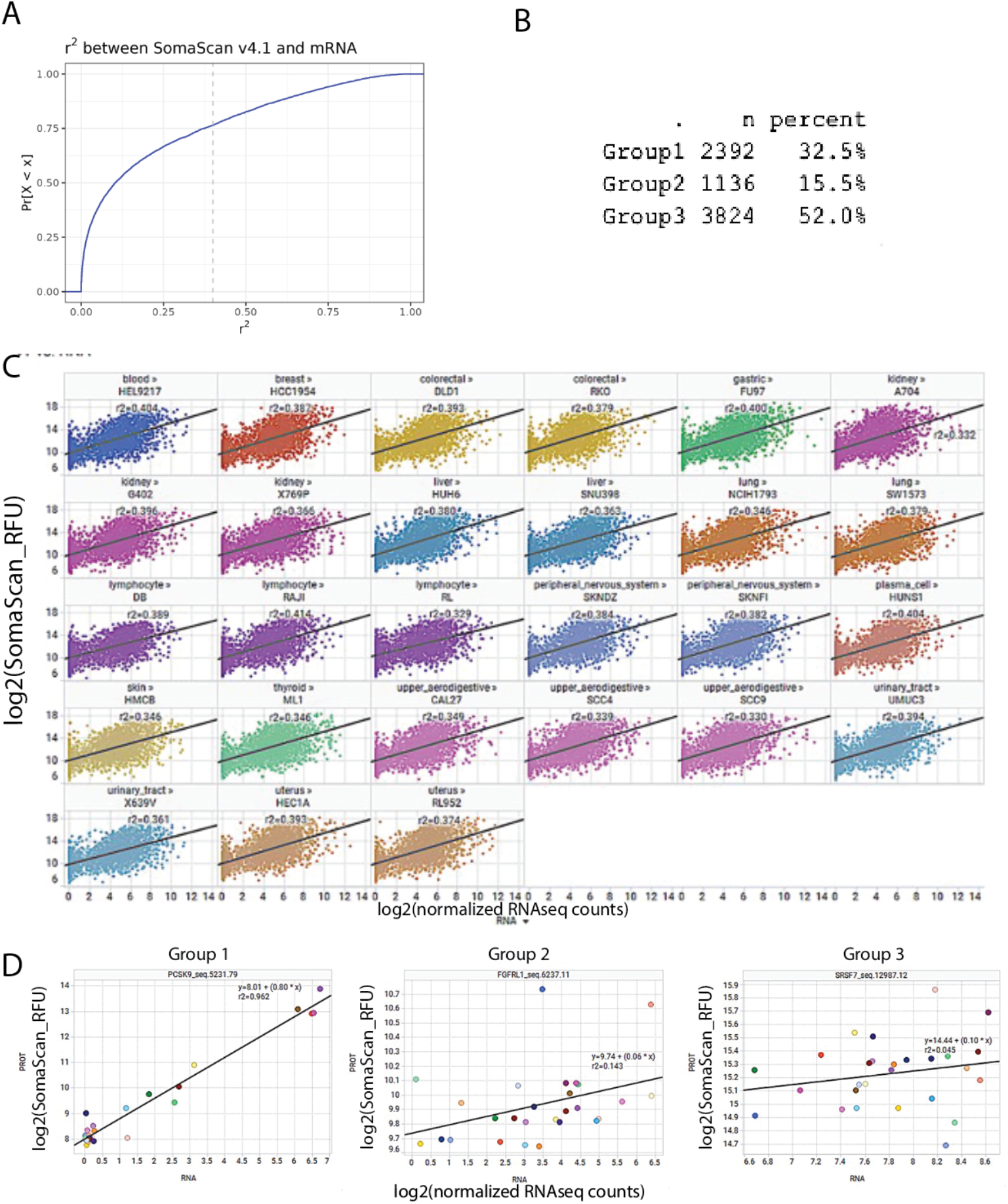
Grouping heuristics evaluating mRNA to protein measurements provide compelling support for approximately one third of the SOMAmer reagents on SomaScan assay. A) Overall correlation of SomaScan measurements to orthogonal mRNA measurements are weak, since they include cases where mRNA data suggest absence of transcripts. B) Table summarizing application of a grouping heuristic to identify cases where there is a range of gene expression levels across the panel provide support for a third of the SOMAmers (Group 1 members). C) This correlation pattern is similar across all cell lines. D) Example SOMAmer reagents by Group designations. Support for endogenous protein binding inference is evident in Group 2 calls.

To characterize SOMAmer reagents by their association to their endogenous mRNA levels, we explored various statistical attributes to explore all the SOMAmer reagents for their potential to measure endogenous protein. Using these parameters, identified groupings for high, intermediate and no confidence in mRNA to protein associate in these data. We found compelling edge cases in certain cell lines, including ones where a single cell line expressed the mRNA and was the only to show an increase in SOMAmer RFU signal. We also noted cases where all cells expressed a narrow range of mRNA expression and there was a corresponding SOMAmer response in a linear model fit (p<0.05). This illustrates the benefits of using a panel of cell lines to establish a correlation between transcriptomic and proteomic data, and contributes to the aggregation of SOMAmer reagent insights described below.

### 3.5. mRNA Correlation Contributes to a Growing Body of Orthogonal Confirmations of SOMAmer Reagent Specificity

The SomaScan platform simultaneously measures more protein targets than any other affinity-reagent technique existing at the time of publication; consequently, the ability to confirm SOMAmer reagents’ specificity to endogenous proteins with orthogonal strategies requires the integration of multiple additional techniques and the rate of orthogonal confirmations lags behind the number of measurements possible with SomaScan. We, along with the scientific community of SomaScan assay users, have approached SOMAmer reagent characterization through a variety of orthogonal methods including antibody techniques (ELISA, proximity extension assays) [24, 25], mass spectroscopy [26, 27], and comparisons to genetic techniques such as cis-protein quantitative trait locus detection (cis-pQTL) analysis [28, 29], and the mRNA correlations reported here (Fig. 5). This represents a current snap-shot of work in progress to provide orthogonal confirmation for the reagents on the SomaScan assay.

**Fig 5.**
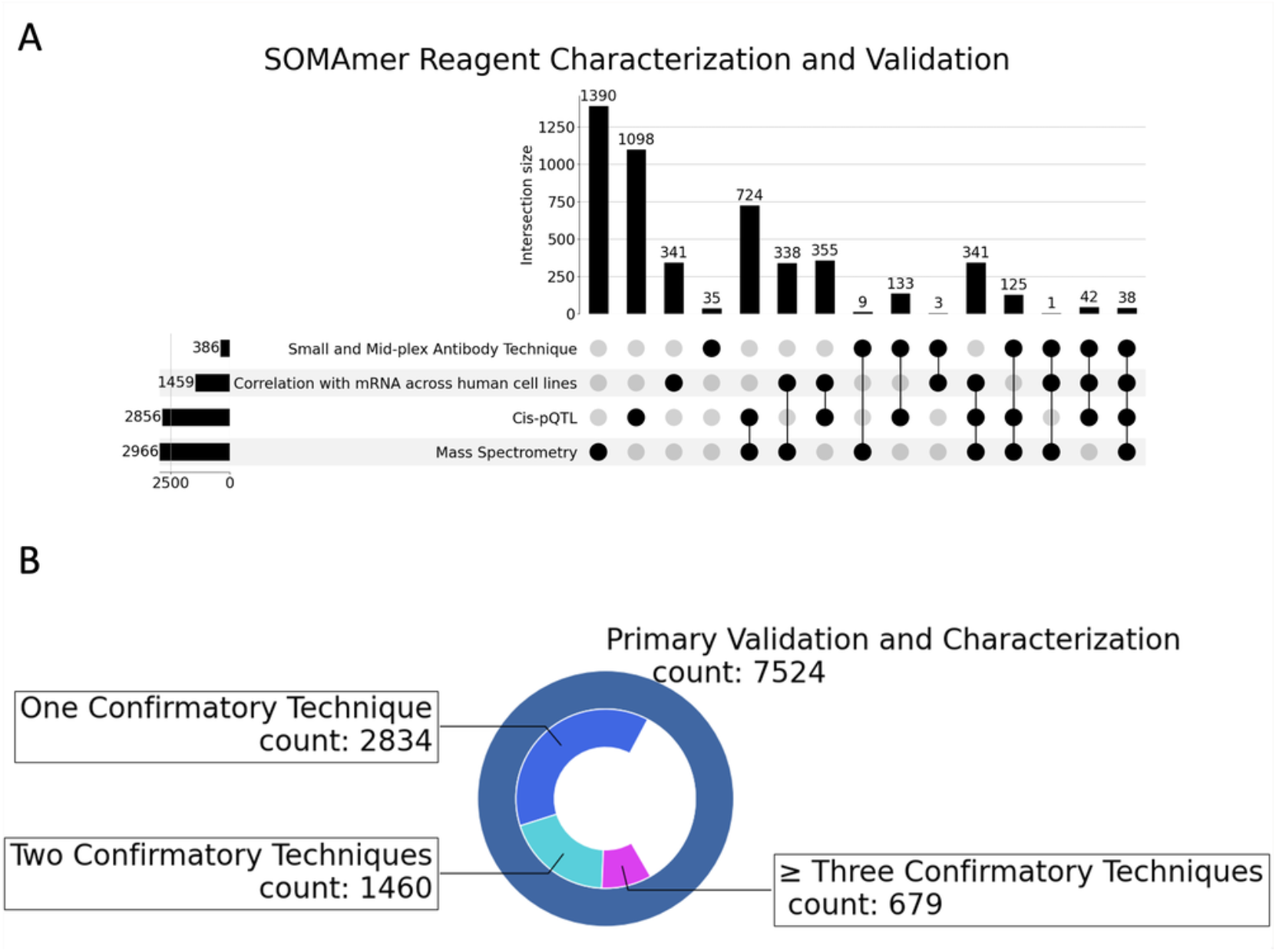
Multiomic integration of evidence for SOMAmer association to its annotated SELEX input protein(s). A) Upset plot of SOMAmer reagent evident for association to endogenous target protein. B) Summary of evidence accumulating for support of SOMAmer reagent binding to the endogenous target corresponding to the SELEX input protein for the SOMAmer reagent.

## 4. Discussion

### 4.1. Application of SomaScan technology to human cell culture advances molecular understanding these models

SomaScan proteome profiling technology permits rapid, multiplex measurement of small sample volumes (< 60 μl per SomaScan run) for biomedical insights. In addition to its primary clinical application measuring blood, SomaScan is suitable for use in other patient sample matrices and in preclinical applications. We demonstrate this here by measuring cell lysates from human cancer cell lines comparing their publicly available steady-state mRNA measurements to the protein measurements on the SomaScan assay. Using multiple unsupervised modeling approaches, we observed clustering of replicate samples and interpretable, higher-order grouping of samples consistent with their known biology.

### 4.1. Topic Modeling provides a mechanism for interpretation of protein subsets within a SomaScan study

Topic modeling has demonstrated an ability to make sense of transcriptomic data and here we demonstrate its use as an approach to interpreting proteomic data at scale. All proteomic samples were clustered by cell line when an unsupervised topic model was applied to SOMAmer-detected protein levels. Even related cell lines of the same cancer type and tissue source were distinguished from one another (e.g. autonomic ganglia neuroblastomas SKNFI and SKNDZ, tongue squamous cell carcinomas SCC4 and SCC9). Related cell lines were often clustered near one another and represented by similar topics, indicating that dimensions spanning multiple cell lines were readily identifiable in the SomaScan assay.

### 4.2. Orthogonal mRNA measurement identifies SOMAmer-endogenous protein association

Following their identification in SELEX experiments, generally with purified recombinant proteins, each SOMAmer reagent included in the SomaScan assay has a unique DNA sequence, unique pattern of diversity-enhancing base modifications, and is synthesized by a fully synthetic solid-phase chemical process. Each reagent is in a sense equivalent to a monoclonal antibody with chemical composition that is precisely defined. This allows for reproducible manufacturing with tight lot-to-lot controls including complete analytical characterization of identity and purity of each reagent, which contributes to high measurement precision as reflected by the exceptionally low coefficients of variance (CVs) of the SOMAscan assay.

Beyond analytical characterization to the original SELEX input protein, it is important to establish that SOMAmer reagents are also capable of recognizing endogenous protein in a biological sample. For that purpose, we used mRNA measurements to set expectation for endogenous protein measurement with SomaScan. By comparing the SomaScan assay to mRNA expression in isolated, well characterized human cancer cell lines, we were able to affirm over three thousand SOMAmer reagents. This provides an opportunity to make robust orthogonal measurements for comparisons for some SOMAmer reagents that may not be possible in other biological matrices, and thereby increase confidence in the performance of both mRNA and proteomic measurement systems. mRNA association in this study have added 341 novel confirmations that were not possible in other matrices and by other measurement systems.

### 4.4. Complementary SOMAmer reagent annotation validation methods expand proteogenomic understanding of the SELEX input protein

To provide independent means of confirming the identity of proteins measured in the SomaScan assay, we have been building a dataset of orthogonal measurements that includes biochemical binding studies with related and unrelated proteins to establish binding specificity and cross-reactivity (which is expected for related proteins that engage SOMAmers with similar epitopes), SOMAmer pull-down experiments coupled with polyacrylamide gel electrophoresis or mass spectrometry [10], genetic associations such as cis-pQTLs and, as described here, correlations with transcriptomic data. Since few of these orthogonal validation tools can individually cover the entire content of reagent in the SomaScan assay, our aim is to use them collectively to provide at least one independent means of ensuring the identity of protein measurements for as large fraction of the SomaScan content as possible.

Recent publications have used correlation to cis- and trans-PQTLs to confirm the measurements made by SOMAmer reagents in samples drawn from human populations [9, 10, 28-39]. Evidence of cis-pQTL associations to a SOMAmer SELEX input protein is interpreted as confirmatory evidence for the SOMAmer binding to its intended, endogenous target. Integrating distinct lines of evidence augments confidence in the performance of the SOMAmer reagents. Taken in concert the UpSet plot above shows that 4973 of the SOMAmer reagents in the SomaScan 4.1 7K menu have been confirmed by correlation to one or more additional techniques. Given that no other proteomic platform measures as many distinct proteoform targets over a wide dynamic range, we expect the number of confirmatory findings to grow from this foundation. The possibility to stimulate cultured cells to modify their gene expression provides fertile ground for future work in this area.

## Supplementary Materials

Table S1: Cancer Cell Line Encyclopedia cell line panel description; Table S2: Topic modeling approach supporting material; Table S3: CCLE panel RNAseq and SomaScan assay normalized data summary; Table S4: mRNA to SOMAmer reagent statistics and association grouping;

## Author Contributions

All authors have read and agreed to the published version of the manuscript. Contributions are: conceptualization, J.L., N.J., L.J. and V.M.; methodology, P.A., F.M., W.S. & J.P.;; formal analysis, D.P., B.W., K.A., T.J. & E.T; writing—original draft preparation, K.D., M.H., A.W., N.J., J.L., D.P., & B.W.

## Funding and Conflict of Interest

This research was resourced as part of a Novartis-Somalogic multiyear collaboration and all authors were employees of Novartis or Somalogic at the time of their contribution to this work.

