## Supplemental_2 for "Application of Cancer Cell Line Encyclopedia for Measuring Correlation Between Transcriptomics and Proteomics as a Guide for System-level Insights"

**SFig 1: Convergence of topic model fit by log-likelihood.** Topic models were fit with the fastTopics R package, with 10 different initializations. The model described in detail in the main text was converged to from two different initial conditions (seeds 5 and 10), whose final fit had the best model log-likelihood (-1.2E08). After initialization, models were fit using expectation maximization for 200 iterations followed by sequential coordinate descent (as described in the fastTopics package documentation) for 800 iterations. Since all models reached convergence by 600 iterations, the figure is truncated along the x-axis from the 1000 total iterations. Note that initial improvements in log-likelihood are considerable, and the figure contains a log-scaled Y axis to highlight convergence in later iterations.

**SFig 2: Samples clustered on topic membership for a 30 topic model.** Within-cell line samples remain clustered, however between-cell line clustering is largely uninformative given that each cell line is captured almost in entirety by one topic.
